# The ultra-sensitive Nodewalk technique identifies stochastic from virtual, population-based enhancer hubs regulating *MYC* in 3D: Implications for the fitness of cancer cells

**DOI:** 10.1101/286583

**Authors:** Noriyuki Sumida, Emmanouil G Sifakis, Barbara A Scholz, Alejandro Fernandez-Woodbridge, Narsis A Kiani, David Gomez-Cabrero, J Peter Svensson, Jesper Tegner, Anita Göndör, Rolf Ohlsson

## Abstract

The relationship between stochastic transcriptional bursts and dynamic 3D chromatin states is not well understood due to poor sensitivity and/or resolution of current chromatin structure-based assays. Consequently, it is not well established if enhancers operate individually and/or in clusters to coordinate gene transcription. In the current study, we introduce Nodewalk, which uniquely combines high sensitivity with high resolution to enable the analysis of chromatin networks in minute input material. The >10,000-fold increase in sensitivity over other many-to-all competing methods uncovered that active chromatin hubs identified in large input material, corresponding to 10 000 cells, flanking the *MYC* locus are primarily virtual. Thus, the close agreement between chromatin interactomes generated from aliquots corresponding to less than 10 cells with randomly re-sampled interactomes, we find that numerous distal enhancers positioned within flanking topologically associating domains (TADs) converge on *MYC* in largely mutually exclusive manners. Moreover, when comparing with several enhancer baits, the assignment of the *MYC* locus as the node with the highest dynamic importance index, indicates that it is *MYC* targeting its enhancers, rather than *vice versa.* Dynamic changes in the configuration of the boundary between TADs flanking *MYC* underlie numerous stochastic encounters with a diverse set of enhancers to depict the plasticity of its transcriptional regulation. Such an arrangement might increase the fitness of the cancer cell by increasing the probability of *MYC* transcription in response to a wide range of environmental cues encountered by the cell during the neoplastic process.

## Introduction

Single cell studies have shown that transcriptional activation occurs in bursts in both prokaryotes and eukaryotes(Sanchez and Golding 2013). The resulting variability in expression levels contributes to transcriptional “noise”, and likely depends on the probability of key limiting events, such as the accessibility to transcription factors and their on-off rates at *cis*-regulatory elements (Hager et al. 2009). *Cis*-regulatory elements, such as enhancers, are often positioned distal to the promoters they regulate - providing yet another level of transcriptional control by 3D chromatin conformation that influences the probability of communication between enhancers and promoters (Fullwood et al. 2009). The arrangement of large domains with enhancing activities, the so-called super-enhancers has been proposed to reduce transcriptional noise and buffer against environmental perturbations in order to robustly maintain the differentiated phenotype (Hnisz et al. 2013; Hay et al. 2016). Super-enhancers can also be activated to drive unscheduled expression of oncogenes, such as *MYC*, and thus the neoplastic process (Hnisz et al. 2013; Loven et al. 2013).

When analyzing the interactomes impinging on enhancers in general and super-enhancers in particular, a common theme is that these interact with each other extensively and that such enhancer hubs collaborate to boost the transcriptional process (Patrinos et al. 2004; Gavrilov and Razin 2008; Berlivet et al. 2013; Kieffer-Kwon et al. 2013; Dowen et al. 2014; Kim et al. 2014; Liu et al. 2014; Markenscoff-Papadimitriou et al. 2014; Xiang et al. 2014; Ing-Simmons et al. 2015). However, experiments attempting to resolve such features have suffered from poor sensitivity or poor resolution of currently available techniques designed to examine higher order chromatin structures. A recurring complication of such techniques, which are all derived from the initial chromosome conformation capture (3C) technique (Dekker et al. 2002), is that they require large amounts of cells to attain resolution. Conversely, although Hi-C protocols (Lieberman-Aiden et al. 2009) have been developed to examine chromatin structure at the single cell level or in small cell populations, they either do not currently have the resolution sufficient to address enhancer-gene communications (Nagano et al. 2013; Stevens et al. 2017) and/or cannot discriminate between real and virtual networks (Nagano et al. 2013; Du et al. 2017; Stevens et al. 2017). Although we have earlier discovered that the 4C technique that we innovated (Zhao et al. 2006) has the capacity to capture multiple interactions to identify instances when several interactions occur simultaneously (Zhao et al. 2006; Sandhu et al. 2009; Gondor et al. 2010; Zhao et al. 2015) to provide important information on chromatin networks using single baits, such information is rare to constitute only a few percent of reads from high throughput analyses, but also requires extensive logarithmic amplification steps. This conclusion is reinforced by our observation that the interactors organizing such networks are rarely clustered together prompting the conclusion that they arise by “dating” rather than “partying” (Sandhu et al. 2009). As there is currently no technique available to address the precise frequencies of enhancer-promoter interactions with the required sensitivity we remain ignorant of their dynamics in small cell populations.

To overcome these limitations, we introduce Nodewalk, which has both the resolution and an unrivalled sensitivity to comprehensively detect stochastic interactions and dynamic changes in chromatin configurations within and between topological-associated domains (TADs)(Dixon et al. 2012) in very small cell populations. We took advantage of the unique features of Nodewalk, such as linear amplification steps, to address the mechanism of enhancer action, to address the mechanism of enhancer action, and examine whether or not simultaneous interactions between enhancers generate cooperating enhancer hubs to drive *MYC* transcription. We document here that although Nodewalk analyses identified a virtual, interconnected core enhancer interactome in relatively large cell populations (corresponding to 10,000 cells), these chromatin structures were largely, if not completely absent in input material corresponding to 21 alleles and by inference at the single cell level. We thus propose that *MYC* interacts with its distal enhancers in a largely mutually exclusive manner. Moreover, the high dynamic importance index of *MYC* in comparison with its enhancers suggests that the position of *MYC* in an inter-TAD boundary region enables it to find and interact with a diverse range of distal enhancers scattered in the flanking TADs, to thereby increase the plasticity of its transcriptional regulation. We discuss how this arrangement might increase the fitness of cancer cells.

## Materials and Methods

### Cell Culture

HCT116 was kindly gifted from Dr. B. Vogelstein and maintained in McCoy’s 5A modified medium (Thermo Fisher Scientific, Waltham, MA. 16600-082) supplemented with 10 % Fetal Bovine Serum (Thermo Fisher Scientific, 10270), penicillin and streptomycin (Thermo Fisher Scientific, 15140122). STR profile was confirmed at Cell Line Authentication Service (LGC standards, Cumberland Foreside, ME) with 77 % match with reference profile. Normal colon cell (HCEC) was obtained from ScienCell, Carlsbad, CA (HCoEpiC, 2950) and maintained in Colonic Epithelial Cell Medium (ScienCell, sc2951). Both cells were cultured under 5 % CO2 circumstance. Drosophila S2 cells were obtained from Thermo Fisher Scientific (R69007) and maintained in Schneider’s Drosophila medium (Thermo Fisher Scientific, 21720024) at the ambient temperature. Cells were routinely screened for mycoplasma contamination using EZ-PCR Mycoplasma Test Kit (Biological Industries, Cromwell, CT. 20-700-20).

### Library preparations of Chromosome Conformation Captured DNA

Chromosome Conformation Capture (3C) was performed as described before(Naumova et al. 2012). Briefly, ten million cells were fixed with 1 % monomeric formaldehyde and the excess amount of the formaldehyde was captured by 0.125 M Glycine. Cells were aliquoted every 1 million and stored at - 80 °C until use. As an external ligation control, equal amount of chromatin from Drosophila S2 cells was fixed in the same way and mixed prior to nuclear isolation. The cells were subsequently collected and suspended in 10 ml of nuclear isolation buffer (0.2 % NP-40, 10 mM Tris-HCl, pH 7.5, 10 mM NaCl, 1x Proteinase inhibitor (Roche, Basel, Switzerland. 04693116001) and incubated for 10 min on ice. Isolated nuclei were precipitated and washed with 1 ml of Buffer 2 (New England Biolabs, Ipswich, MA. B7002S). Next, nuclei were re-suspended in 500 μl of 1.2x buffer 2 and un-crosslinked fraction was removed by adding 5 μl of 10 % SDS and the incubation for 1 hr at 37 °C. Excess amount of SDS was captured by Triton X-100. The chromatin fiber was fragmented by the incubation at 37 °C overnight with *Hin*d III (1 U/μl, New England Biolabs, R0104T). The fragmentation was stopped by the incubation at 65 °C for 20 min with 1 % SDS. 80 μl of the solution was transferred to a fresh tube and saved for checking the digestion (Additional file 1: Fig. S1A). The residual solution was transferred to a 50 ml tube containing 7 ml of ligation buffer (50 mM Tris-Cl, pH 7.5, 10 mM MgCl2, 1 % Triton X-100) and incubated for 1 hr at 37 °C followed by the addition of 70 μl of 100 mM ATP, 70 μl of 1 M DTT and 2000 U of T4 DNA ligase (New England Biolabs, M0202M). The solution was incubated at 16 °C overnight followed by 1 hr at RT. For the reversal of the crosslinks, the solution was incubated with 300 μl of 5 M NaCl, 350 μl of 10 % SDS and 50 μl of 10 mg/ml Proteinase K at 65 °C for at least 4 hrs. Finally, DNA was collected by conventional ethanol precipitation followed by the purification of ligated DNA with MSB Spin PCRapace (Stratec, Birkenfeld, Germany. 1020220200) and Zymo Clean & Concentrator-5 (Zymo Research, Irvine, CA. D4013). The quality of 3C DNA was visualized on 1 % agarose gel (Additional file 1: Fig. S1A,B)(Naumova et al. 2012).

### Adaptation of 3C protocol for low cell input

The 3C libraries were prepared as above with following modifications (see Additional file 1: Fig. S3 for additional information): Following formaldehyde fixation, HCT116 and S2 cells were counted and diluted in nuclear isolation buffer to make 600 cells/μl (corresponding to ∼3 ng of genomic DNA/μl). Aliquots (0.5 μl) of the resulting cell suspension was mixed and incubated on ice for 10 min. The cell suspension was directly diluted 10 times with x 1.2 Buffer 2 (see above). Next, the samples were digested with *Hin*d III and ligated as above, albeit in a smaller reaction volume (20 μl for *Hin*d III digestion; 200 μl for 3C ligation). Following reversal of the crosslink, the 3C-DNA was purified using the ChIP DNA Clean and Concentrator kit (Zymo Research, D5205). The elution buffer was pre-heated at 65 °C to increase the recovery of large DNA fragments.

### Measurement of restriction digestion efficiency

The digestion efficiency was estimated by qPCR (Additional file 3: Table 2) (Gondor et al. 2008). To evaluate the digestion efficiency of crosslinked chromatin we designed F1/R1 and F2/R2 flanking the *Hind* III sites at the 5’ and 3’ends of the *MYC* promoter and gene body. The amount of total DNA was quantified with the primers F3/R3 used to produce a PCR fragment lacking an internal *Hind* III site. The linear range of amplification was determined by serial dilution of sonicated genomic DNA. The digestion efficiency was calculated as (1-(PCR_F1+R1/F2+R2_/PCR_F3+R3_)) × 100 (%).

### 3C-qPCR

To quantify the frequency of 3C ligation, we first generated control 3C products. Briefly, naked genome DNAs were isolated from either HCEC or HCT116 using Mammalian Genomic DNA Extraction Kit (Sigma Aldrich, St Louis, MD. G1N10). Then genomic regions containing the annealing site of 3C primers and neighboring *Hin*d III site were amplified by PCR. Each amplicon was digested with *Hin*d III, pooled into one tube and ligated. Next, chimeric DNA of bait and each interactor were amplified by 3C primers (Additional file 3: Table S2). The correct amplicons were recovered from agarose gels and quantified by Nanodrop ND1000 (Thermo Fisher Scientific). The linearity and the sensitivity of 3C qPCR was confirmed using the serial dilution of the control 3C products mixed with sonicated genome DNA (200 ng per reaction). QPCRs were performed using 200 ng of 3C library by referring the same dilution of control 3C products.

### Nodewalk analyses

3C library was tagmented using either Nextera DNA sample prep kit for routine or Nextera XT DNA sample prep kit for the analyses with lower input (Illumina, San Diego, CA. FC-121-1031, FC-131-1024). Amount of the input was validated by qPCR (Additional file 3: Table S2). Primer 1a and 1b (Fig. 1A and Additional file 5: Table S4) were incorporated by limited cycle PCR following the manufacturer’s instruction with replacing Nextera primers with 22 pmol each of Primer 1a and 1b. For Nodewalk with 34.8 pg of input, PCR cycle was increased to 17 following the previous research (Picelli et al. 2014). The library was amplified by *in vitro* transcription using Maxiscript T7 kit (Thermo Fisher Scientific, AM1312M). The template DNA was removed by Turbo DNase I (Life technology, AM2239) prior to the RNA purification step (RNeasy, Qiagen, Venlo, Netherlands. 74104). For primer annealing, 1 μg of template RNA and 10 pmol of bait-specific primer (primer 3a, Fig. 1A) in 20 μl of water was denatured at 75 °C for 3 min and then mixed with 25 μl of 2x buffer of Platinum Quantitative RT-PCR ThermoScript One-Step System (Thermo Fisher Scientific, 11731-015) and 1 μl of SUPERase In RNase inhibitor (Thermo Fisher Scientific, AM2694). The solution was incubated at 4 °C for 44-48 hrs with presence or absence of betaine (see following section). Following the incubation at 65 °C for 5 min, the primer was extended by adding 1 μl of Thermoscript Plus/Platinum Taq mix at 60 °C for 30 min. The enzyme was inactivated by incubation at 95 °C for 5 min, and then the bait-interactor chimeric DNA was enriched by amplification with 1 μl of Illumina N5 primer from Nextera DNA sample prep kit using the following PCR cycle: 36 cycles of 95 °C for 10 sec, 63 °C for 30 sec and 72 °C for 30 sec followed by final extension at 72 °C for 3 min. The resulting library DNAs were purified by same volume of Agencourt Ampure XP (Beckman Coulter, Brea, CA. A63880). To extrapolate the specifically primed 3C cDNA, one tenth of the libraries were digested with *Hin*dIII for 1 hr at 37 °C and subjected on High Sensitivity DNA Analysis Kit (Agilent technologies, Santa Clara, CA. 5067-4626) with Bioanalyzer 2100. Only the library that showed the single band corresponds to the enriched bait sequence was used for the high throughput sequencing (Fig. 1D). Finally, Illumina P7 sequence was incorporated by 5 cycles of PCR with Primers 3a and 3b and the resulting Nodewalk libraries were purified by using the same volume of Agencourt Ampure XP. Maximum 10 libraries were pooled and sequenced on Illumina Miseq (Illumina) using Miseq reagent cartridge v2 (Illumina) that generated 140-150 bp paired-end reads.

**Figure 1:**
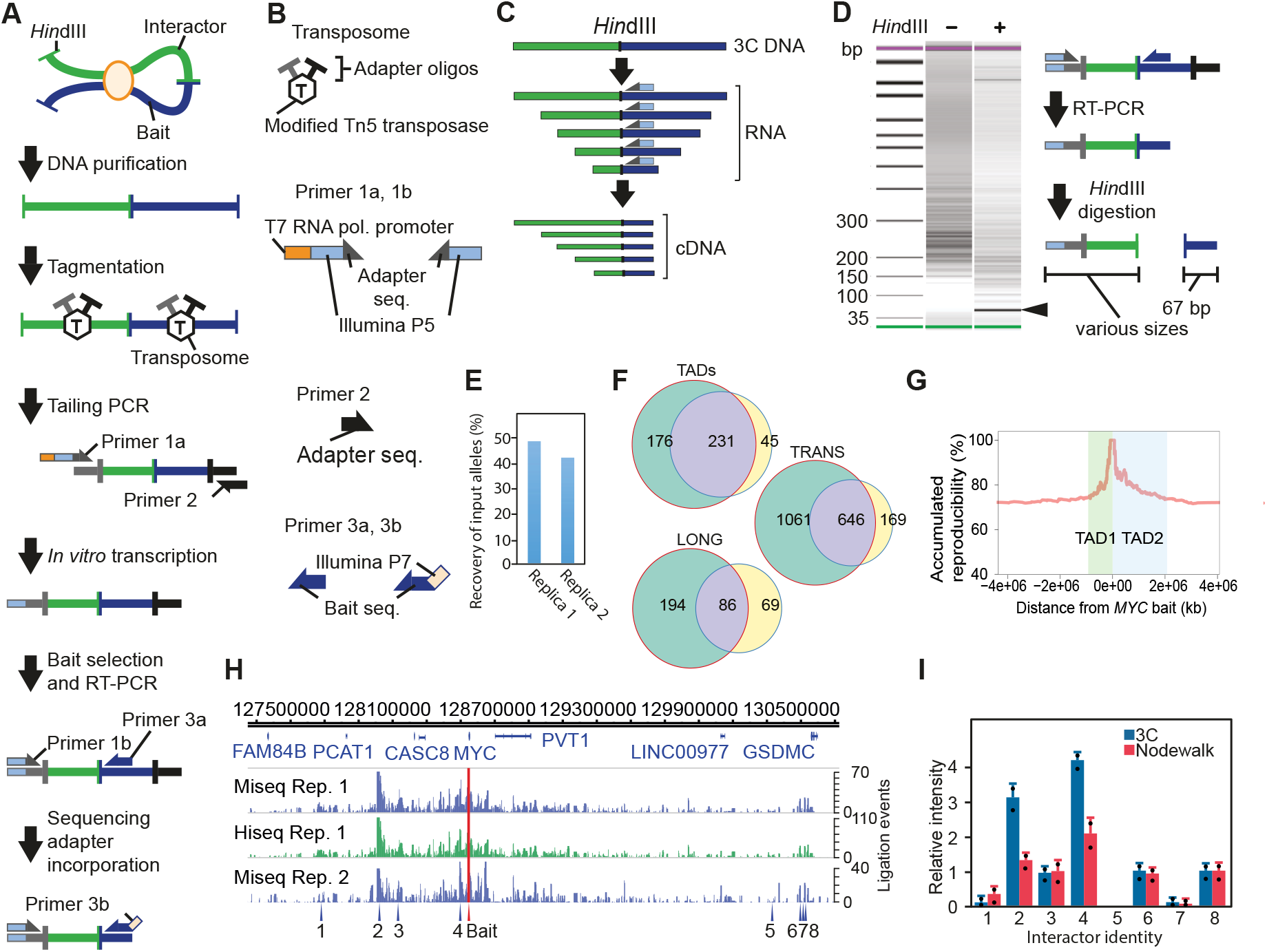
The Nodewalk principle and associated quality controls. A) Schematic representation of Nodewalk procedure. Region of the interest (Bait: blue) and interacting locus (Interactor: green) are represented by line with restriction site (*Hin*d III). Horizontal arrows indicate primers. B) Schematic representation of oligo DNAs and primers designed for the Nodewalk protocol. C) Principle to generate cDNAs from 3C RNA. The size of the bait becomes uniform after the enrichment of the bait-interactor junction. D) Assay to evaluate the fold enrichment of specifically primed 3C cDNAs. The DNA band indicated by an arrow represents the enriched bait fragment. Panel E) shows the recovery of interactors between two independent replicates while F) shows the amount of reproducible interactors between two independent replicates stratified as indicated in the panel. G) Accumulated reproducibility between two independent experiments. H) Map of *MYC* locus with arrows indicating the position of interactors identified by using *MYC* as initial bait. I) Comparison between Q-PCR analysis of 3C DNA products and the resulting normalized reads from the very same sample. Data are represented as mean + SEM from two independent replicates. Dots indicate the actual values. The numbers indicate the positions of interactors identified from the Nodewalk analysis shown in H).

### Sampling of low input control

For Nodewalk analysis of 10 aliquots of 0.88 ng of 3C DNA were sampled from 3C DNA prepared from 1 million cells. To assess technical variation, three aliquots of 1 μg each of 3C RNA were subjected to Nodewalk analysis. Aliquots of 34.8 pg of 3C DNA (corresponding to genomic DNA of 7 cells which contains 21 *MYC* alleles) were sampled from a diluted library of one of the 3C DNA prepared from 300 cells.

### Optimization of primer annealing

Following the preparation of the 3C library by a conventional protocol, 3C RNA was generated by *in vitro* transcription (see above). The bait primers were annealed at 4°C irrespective of its estimated melting temperature with the stringency of the annealing process provided by betaine at either 0.5, 1 or 1.5 M (final conc.). The betaine concentrations of each primer used here are summarized in Additional file 5: Table S4.

### Sequence mapping and filtering

Paired-end reads were independently mapped using bwa version 0.7.12-r1039 to a merged reference genome composed of phiX174, Drosophila (BDGP5.65), Escherichia coli K12 and human (GRCh37.75) genomes. Resulting SAM files were processed and annotated with the Nodewalk Pipeline Analysis software (Fernandez-Woodbridge and Sumida, manuscript in preparation). Briefly, alignments with mapping quality greater than 10 of the second read were used to determine probe position. An extension region (extending from the probe’s end to the first restriction site) was used to discriminate valid from mis-annealing events. The total number of alignments in the first read with the proper probe extension in the second read was reported by restriction fragment.

### Window summaries at Nodewalk with low input

Interactors detected by Nodewalk with lower input such as 34.8 pg or 0.88 ng of 3C DNA or 177 cells aliquots were summarized in 10 kbps window. At first the reference genome was divided into 10 kbps window and each interaction was binned according to the center of the interacting locus. LEs were summed when more than one interactor was overlapping with a window.

### Identification of statistically significant interactions

To determine the statistically significant interactions, the empirical resampling approach for determining significant enrichment described in reference (Williams et al. 2014) was adopted. In brief, the observed reads on each chromosome were randomly associated (shuffled) with a *Hind* III fragment from that chromosome. Next, a significance threshold, the minimum number of reads (X) in each *Hind* III fragment required to achieve a false discovery threshold (FDR) lower than 0.01, was calculated, i. e.:

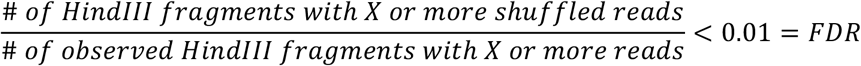

We repeated the shuffling for 1,000 times to obtain a distribution of X at each chromosome. All of the analyses calculated a significance threshold from the top fifth percentile. Because regions around the baits are expected to have a higher background, the regions composing single peaks around the bait were omitted. The above approach was applied to all Nodewalk samples except the libraries with 21 alleles, due to small sample limitations. Finally, to avoid possible PCR biases, the number of distinct transposon integration sites was reported as output (ligation events – or LE).

### Enrichment analysis of cis regulatory elements

The number of interactors with each overlapping ChIP-seq peaks (Hits) was compared to a random model through a permutation based approach by running over 1,000 random shuffles. At each random iteration, we generated a library of interactors and counted the number of peaks overlapping the ChIP-seq library (rHits). The random library maintained the same number of long (defined as interactors are >5 Mbps flanking genomic distance from the bait), short (<5 Mbps), local (<100 kbps) and *trans* interactors as the observed library. The permuted p-value is reported as the number of times in which the rHits was higher than the observed Hits divided by the total number of permutations. ChIP-seq peaks were integrated on the lists of the interactors using “findOverlaps” function from “GRanges” R package with setting of maxgap = 0. Reference ChIP-seq data are listed in Table 1.

**Table 1.** External Datasets.

| Roadmap Epigenomics |  |  |
| --- | --- | --- |
| Colonic Mucosa H3K4me1 | GSM621670 | NarrowPeak for E075<br>http://egg2.wustl.edu/roadmap/data/byFileType/peaks/consolidated/narrowPeak/<br>Note: Substitute for HCEC |
| Colonic Mucosa H3K27ac | GSM1112779 |  |
| Geo Entries |  |  |
| HCT116 H3K4me3 | GSM1240115 | GSE51176: HU Dataset: Mapping: mapped to HG19 with bwa 0.7.12 @ MapQ>10, Peak Calling: macs 2.0.9 (Params: -g hs -p 1e-2 -nomodel -c GSM1240117) |
| HCT116 H3K4me1 | GSM1240111 |  |
| HCT116 H3K27ac | GSM946854 | GSE38477: Cell type-specific binding patterns reveal that TCF7L2 can be tethered to the genome by association with GATA3. Analysis: Mapping: mapped to HG19 with bwa 0.7.12 @ MapQ>10. Peak Calling: macs 2.0.9 (Params: -g hs -p 1e-2 -nomodel -c GSM1240117) |
| ENCODE |  |  |
| HCT116 CTCF | GSM1022652 | GSE30263: CTCF Binding Sites by ChIP-seq from ENCODE/University of Washington |
| HCT116 Rad21 | GSM1010848 | GSE32465: Transcription Factor Binding Sites by ChIP-seq from ENCODE/HAIB<br>Analysis: mapped to HG19 with bwa 0.7.12 @ MapQ>10, macs 1.4 no control |
| Other |  |  |
| HCT116 Super Enhancer | Reference(Hnisz et al. 2013): Data S1, filtered only SuperEnh = 1 |  |
| Colon Crypt Rep1 (HCEC stand-in) Super Enhancer | Reference(Hnisz et al. 2013): Data S1, filtered only SuperEnh = 1, Note: Substitute for HCEC |  |
| HG19 Ensemble Genes 75 | http://feb2014.archive.ensembl.org/biomart/martview/aabe8088059e12c608bc3805af31b156 |  |
| Drosophila Genome | BDGP5.65 |  |
| Phix Genome | gi\|9626372\|ref\|NC_001422.1\| Enterobacteria phage phiX174, complete genome |  |
| E. coli Genome | K12 |  |
| Human Genome | GRCh37 |  |

### Reduction of LE for the comparison of the enhancer proportion between HCEC and HCT116

Since there are 3 *MYC* alleles in HCT116, we selected 2/3 of the interactions in HCT116 based on the Probability 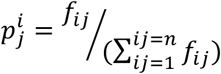 where *f_ij_* is the frequnecy of interaction between node i and node j and n is number of interactions (LE). This calculation reduced the total interactors in HCT116 from 10141 to 7250.

### Calculation of the recovery of ligation events and quantification of the input cells

The recovery of the ligation events was defined as the amount of all observed detected ligation events compared to the expected amount of all possible ligation events using the number of the de-duplicated reads (LE). For this calculation, we included LE of non-significant interactions. The sum of LE was therefore divided by the number of the possible LE estimated as follows: Using the representative amount of genomic DNA in human cells (5 pg/cell), we estimated the number of the input cell from the quantity of 3C DNA estimated by either Nanodrop (for the input of 10,000 cells) or qPCR (for the experiments with lower input, see above and Additional file 3: Table S2). As HCT116 cells harbor 3 *MYC* alleles, the estimated number of alleles per cell was therefore multiplied with 3 for HCT116 and 2 for HCEC. These values indicate the potential of the ligation events at the one side of the restriction site in the bait fragment, including self-ligation as well as interactions with neighboring fragments. As the overall recovery of ligation events exceeded 100 % at Nodewalk with 34.8 pg input, caused by the addition of non-template base or nibbling at the 3’-end of the interactor generated by higher number of PCR cycles, LE exceeding 1 were counted as 1.

### Mapping the interactors with increased window size

Ligation events of 9 Nodewalk libraries derived from 300 cells or 10 libraries derived from 0.88 ng of 3C DNA were summed up at each window size (either 25, 50 or 100 kbps). The windows were shifted each 10 kbps and visualized on WashU Epigenome Browser (Zhou et al. 2011) (summary method: Average).

### Comparison between expected and observed number of enhancers within the MYC interactome

A permutation-based approach was adopted to test whether the observed number of enhancers impinging on *MYC* in the low cell input protocols, namely the Nodewalk with 0.88 ng and with 34.8 pg inputs, significantly differs from the corresponding expected numbers. Specifically, a null distribution was formed by randomly selecting interacting regions 10,000 times from the *MYC* network with 10k cells input keeping the numbers of regions as of the corresponding observed low input interactomes. The method assigned the sampled interactors with probability weights which were based on their observed number of ligation events (LE) in the higher cell count network. In each cycle, the total number of interacting regions that overlapped with H3K27ac peaks was counted. The approach was repeated separately for each analysis.

### RNA/DNA FISH analyses

Single stranded probes were prepared to increase the efficiency of the annealing. At first double stranded DNA spanning *MYC* intron 1 (chr8:128,749,271-128,750,480) and a part of intron 2 (chr8:128,751,280-128,752,201) were obtained by PCR (5’ terminal of forward primer is biotinylated to capture the sense strand, see additional file 3: Table S2). DNAs were labeled with Green 496 dUTP (Enzo Life Sciences, Farmingdale, NY. ENZ-42381) for intron 1 and Chroma Tide Texas Red −12-dUTP (Thermo Fisher Scientific, C-7631) for intron 2 using Bioprime Array CGH kit (Life technologies, 18095-011) and captured by Dynabeads M280 Streptavidine (Thermo Fisher Scientific, 11205D). The beads were incubated at 98 °C for 5 min and released anti-sense strand was recovered. At last single stranded DNA was purified with Zymo Clean & concentrator −5. RNA FISH analyses were performed as previously described(Zhao et al. 2015). In brief, cells cultured on chamber slides (Thermo Fisher Scientific, 154534) were crosslinked with 3 % formaldehyde and stored in 70 % Ethanol at −20 °C until use. In all steps the ribonuclease inhibitor Ribonucleoside Vanadyl Complex (NEB, S1402S) was added to the buffers. Cells were rehydrated in PBS, permeabilized with 0.5 % Triton X-100 in 2 x sodium salt citrate (SSC) for 10 min at RT. The FISH probe mixed with human Cot-1 DNA (Thermo Fisher Scientific, 15279-011) was hybridized with 2 x SSC, 50 % formamide and 10 % dextran sulfate overnight at 37°C. Cells were washed twice with 2 x SSC/ 50 % formamide for 15 min at 40 °C and 2 x SSC for 15 min at 40 °C and mounted with Vectashield mounting medium with 4,6-diamidino-2-phenylindole (DAPI) (Vector Labs, Burlingame, CA. H-1200). DNA FISH was performed as same as RNA FISH but with denaturing the genome DNA in 2 x SSC/50 % formamide for 40 min at 80 °C prior to anneal the probes.

### ChrISP analyses

The ChrISP probe was prepared from a pool of 4 PCR products spanning the *MYC* promoter and its gene body (chr8:128,746,000-128,756,177), a pool of 2 PCR products spanning the part of CRC super-enhancer (E, chr8:128,216,526-128,225,855) harboring highest LE with bait 5 at HCT116 and a pool of 2 PCR products spanning the negative control site (chr8:127,220,074-127,229,883) where bait 5 was not interacting with. The PCR products were sonicated and labeled with Digoxigenin-11-dUTP (Roche, 11573152910) or Biotin-16-dUTP (Roche, 11093070910) using Bioprime Array CGH. Formaldehyde-fixed (1%) cells were hybridized with ChrISP probes as described in RNA/DNA FISH protocol. Next, the cell slides were washed twice with 2 x SSC/ 50 % formamide for 15 min at 40 °C and 2 x SSC for 15 min at 40 °C. After 1 hour blocking at RT cells were incubated with anti-biotin antibody (Abcam, Cambridge, UK. ab53494) and anti-Digoxigenin (Roche, 11333062910) overnight at 4 °C. The following ChrISP assay was performed as previously described(Chen et al. 2014a; Chen et al. 2014b) and the ChrISP probes were visualized by using the secondary fluorescent antibodies (goat anti-rabbit IgG Dylight 594, 35560, Thermo Fisher Scientific; goat anti-mouse IgG Dylight 550, 84540, Thermo Fisher Scientific) for biotin and digoxigenin, washed again with PBS/ 0.05% Tween20 and mounted with Vectashield mounting medium with 4’,6-diamidino-2-phenylindole (DAPI). As a technical control, a sample omitting the mouse secondary antibody was included in parallel, called R plus control, in each experiment. False discovery rate (FDR) was defined as the number of ChrISP signal at test sample/R plus control at each threshold.

### Grid confocal microscopy

Cell imaging and generation of optical section in 3D were carried out on Leica DMI 3000B fluorescent microscopy with OptiGrid device (Grid confocal) using Volocity software (Perkin Elmer, Waltham, MA). Stacks were taken at 0.3 μm intervals in the Z-axis. In each experiment 150-300 alleles were counted (Additional file 7: Table S6).

### Network visualization

All the network figures were generated by Gephi 0.9.1 (Bastian M. 2009) (layout: Force Atlas 2).

### Bootstrapping network generation

From the reads computed after Nodewalk data processing we generated a set of networks that represent the family of possible networks that can be originated from the sequencing data by bootstrapping. To this end, we first selected a number of reads to be included (n=500, 1,500, 5,000) and randomly selected with replacement a number of reads equal to n from the pool of reads. The number of bootstrapped networks computed was 1,000. We then calculated 95% CI (Confidence Interval) of the median of influence index and entropy on these 1,000 weighted network (weights are number of reads) using bootstrap technique. There is no overlap between CIs and the values of both measures; both influence index and entropy are significantly different in HCT116 and HCEC network. (Both median and mean p < 2.2e-16 using Wilcoxon and, Welch’s t-test respectively).

### Dynamic importance index

We used Eigenvalue centrality that quantifies the role of each node in propagating signal through the whole network to estimate the node influence metric using igraph package (Csardi 2006) of R.

## Results

### The Nodewalk principle

The Nodewalk method is built on the initial 3C technique (Dekker et al. 2002), but with several key modifications (Fig. 1A). To ensure optimal and reproducible results, cells were cross-linked with freshly prepared 1% formaldehyde (FA) solution, instead of formalin (Gondor et al. 2008). The crosslinked chromatin DNA was digested to near completion (on average 92%) with *Hind* III while minimizing its star activity and ligated to completion (Additional file 1: Fig. S1A). Following de-crosslinking and DNA purification, a modified Tn5 transposase introduced random DNA cuts, which enabled the elimination of PCR duplicates (see below) and ligated the adaptor oligos (Fig. 1B). This step generated uniform and small fragment sizes of 150-250 bp (Additional file 1: Fig. S1B). The T7 RNA polymerase promoter sequence flanking Illumina P5 sequence was incorporated by tailed PCR (5-17 cycles depending on the size of the input material). To capture the 3C products, the ligated genomic DNA was converted to RNA, enabling a linear, 1,000-fold amplification of 3C sequences, followed by reverse transcription using primers positioned close to the restriction enzyme site of the region of choice (Fig. 1A). Finally, Illumina sequence adapters were incorporated using the same primers equipped with P7 sequence against the cDNA that already contained P5 sequences (Fig. 1A,B). This arrangement enabled the direct generation of double-stranded cDNA, suitable for sequencing. Spurious ligation events were routinely assessed by spiking human chromatin with similarly cross-linked and digested chromatin derived from Drosophila S2 cells. Following high throughput sequencing, this strategy demonstrated that the proportion of human-Drosophila chimeric reads never exceeded 1%.

As initial bait, we chose *MYC* as this region is not only a key to understand the neoplastic process, but has already been used in 3C (Xiang et al. 2014) and other 3C-derived analyses (Li et al. 2012; Shi et al. 2013; Dryden et al. 2014; Rao et al. 2014). The estimate of the enrichment of sequences specific for the bait (Fig. 1C, D) was based on that each *Hind* III fragment represents on average one 830,000^th^ of the human genome. As the specifically primed bait cDNA corresponded to 50 +/-10 % of the entire sequence population, the strategy to prime the 3C RNA library with dedicated oligos achieved >400,000-fold enrichment of the bait and its interactors in one single step (Fig. 1D). To remove PCR duplicates which is essential for quantitative analysis (Schwartzman et al. 2016), we assessed the number of the cutting sites generated by the transposase, hereafter termed “ligation events or LEs” for counting each unique interaction, rather than the raw reads in the sequence library (Additional file 1: Fig. S1C). Comparing two independent replicas, Nodewalk was able to recover ca 45% of all bait alleles (Fig. 1E) with a reproducibility of interacting sequences highest within the neighboring TADs (Fig. 1F,G). To validate the *MYC* interactome, we focused on 8 different regions representing high and low-abundant interactors in *cis* (Fig. 1H, Additional file 2: Table S1, Additional file 3: S2). Conventional quantitative 3C analysis confirmed only minor biases when compared with the number of de-duplicated reads (LE) obtained by high throughput sequencing (Fig. 1I). We conclude that the Nodewalk strategy correctly measured the relative frequencies of chromatin fibre interactions with a high reproducibility and recovery from 3C DNA aliquots representing only 10,000 cells. As there was no discernible difference in data output when the same sample was analyzed by either Mi-seq and Hi-seq (Fig. 1H), we also conclude that the Nodewalk technique is cost-effective.

### Walking the nodes: A network of enhancers

The inclusion of an *in vitro* transcription step in the Nodewalk protocol represents a unique advantage in comparison with competing techniques as the RNA template can be reproduced numerous times from the same initial library, offering the possibility to establish the interactomes of multiple interactors in a sequential manner (Fig. 2A) even if the initial sample size is small. The statistically significant interactors were selected using a background algorithm established by (Williams et al. 2014) (see Methods). The implementation of this principle is shown for the *MYC* locus (Fig. 2B), with 9 central nodes enriched in enhancer marks (Fig. 2C,D) selected from iterative screens (Additional file 4: Table S3, Additional file 5: S4). We conclude from these analyses that each of the selected baits reconnected with the original bait to further document the reproducibility of the network. Moreover, the network includes frequent inter-chromosomal interactions (Additional file 1: Fig. S2A) and displays enrichment for enhancer marks (Additional file 1: Fig. S2B) both in *cis* and in *trans*, suggesting that active domains functioned as preferred interacting partners.

**Figure 2.**
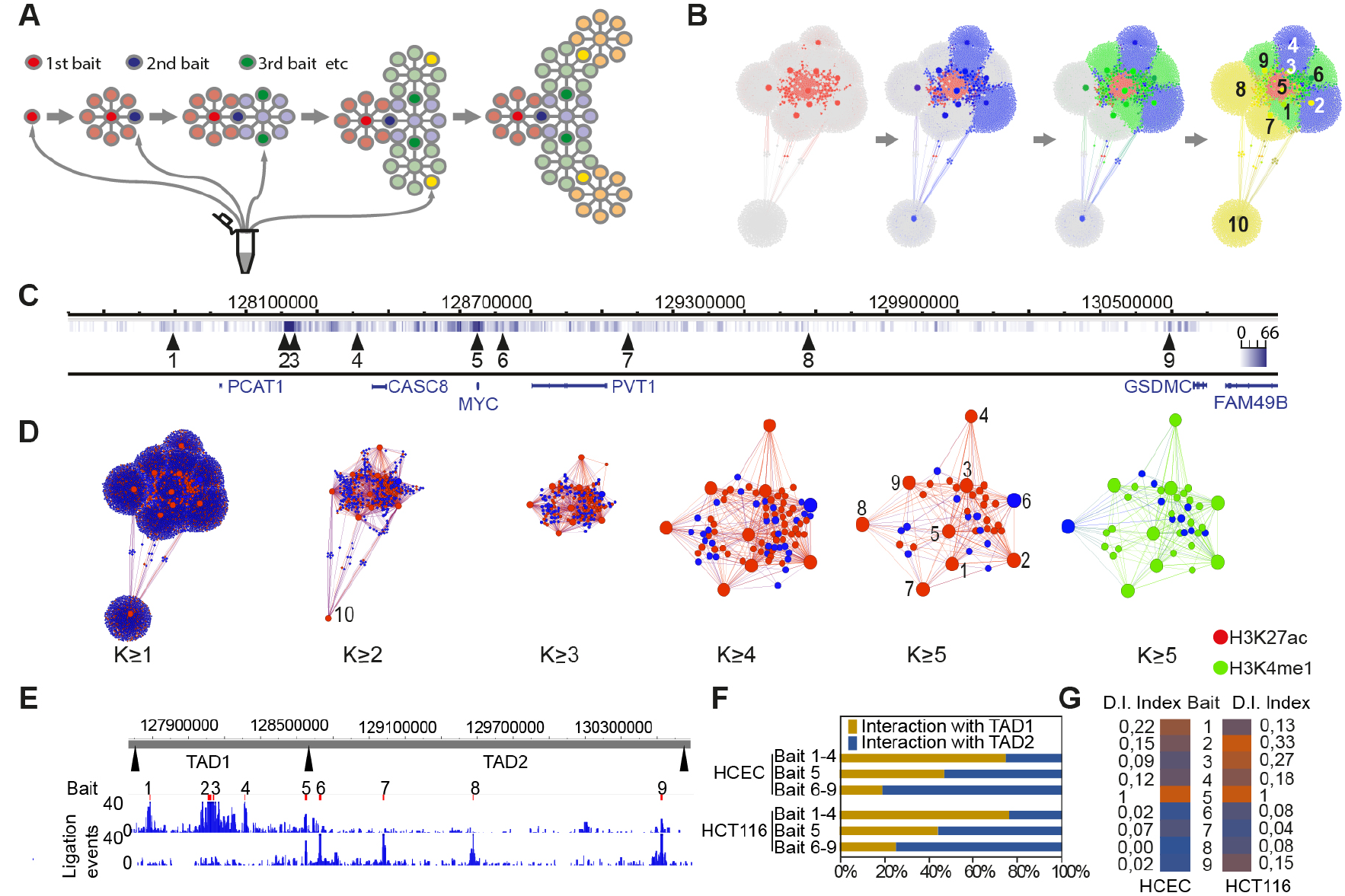
The generation of Nodewalk networks and their link to enhancers. A) Schematic visualization of sequential “Nodewalking”. The iterative nature of the principle is represented by a network of detected loci (circles) and their interactions (lines). B) The actual network generated from the *MYC* locus in HCT116 cells, using this strategy. C) The position of each new bait highlighted in B). All baits were from *MYC* and the TADs flanking *MYC* on chromosome 8 except for bait nr 10 that originated from chromosome 5. Vertical lines indicate the interactors and their ligation events (LE) impinging on *MYC*. D) The network structure from HCT116 cells stratified by its k-core values. The red and green nodes identify regions overlapping with H3K27ac and H3K4me1 peaks, respectively. The size of each node reflects the number of the interactors. E) Distribution of interactors generated from enhancer baits from within TAD 1 and 2, respectively. F) The interactions of enhancer hubs largely follow the TAD boundaries - with the exception for the *MYC* bait (nr 5), which equally interacts with both flanking TADs in both HCEC and HCT116 cells. G) Dynamic importance index (D.I. Index) analysis of the nine enhancer baits in HCEC and HCT116 cells.

To better understand the underlying dynamics, the networks from HCT116 cells and their normal counterparts, primary colon epithelial cells (HCEC), were stratified according to their k-core values, which measure the cohesion of the network (Seidman 1983) (Fig. 2D and Additional file 1: Fig. S2C). To compensate for any bias in network comparisons arising from the fact that *MYC* is triploid in HCT116 (Langer et al. 2005) and diploid in HCEC cells, we randomly sampled two thirds of the interactions from the HCT116 network. Although the un-stratified network displayed only minor differences between the normal and cancerous counterparts, the most connected nodes were considerably more prominent in colon cancer cells than in normal cells (Additional file 1: Fig. S2C,D). Examining the chromatin marks associated with higher connectivity, we found that primed or active enhancers were strongly enriched with increasing k-core values specifically in cancer cells (Additional file 1: Fig. S2C,D). Although this suggested that active enhancer-specific histone modifications serve to increase network connectivity, a fraction of the well-connected chromatin hubs present in HCT116 cells interacted also in HCEC cells despite not carrying H3K27ac marks in these cells (Additional file 1: Fig. S2C-E). Finally, as could be expected, the network nodes displaying the highest connectivity preferentially localized within the two TADs flanking *MYC* (Fig. 2E, F). Interestingly, *MYC* itself is located close to the inter-TAD boundary and is able to explore the neighboring two TADs equally well in both HCEC and HCT116 cells (Fig. 2F). This conclusion is in keeping with the observation that *MYC* has the highest dynamic importance index within the chromatin interactome for both HCEC and HCT116 cells (Fig. 2G). The dynamic importance index represents the importance of the node in keeping the network structure, i.e. removing a node with high index will change the structure of the network significantly and globally in contrast of some other centrality indexes such as the degree which are local indexes (Lohmann et al. 2010). Intuitively, the importance of a node does not only depend on the number of neighbors it has, but also how central its neighbors are.

### Virtual enhancer hubs emerge from stochastic enhancer-promoter communications in large cell population

The presence of an enhancer hub organized around an active gene and involving primarily local sequences distributed in the neighboring TADs is in keeping with numerous published observations (Patrinos et al. 2004; Gavrilov and Razin 2008; Kieffer-Kwon et al. 2013; Dowen et al. 2014; Dryden et al. 2014; Hughes et al. 2014; Markenscoff-Papadimitriou et al. 2014; Ji et al. 2015) using assays that rely on large cell populations as input material. To understand the mechanism of enhancer action in 3D and the plasticity of transcriptional regulation in development and disease, we set out to explore whether or not multiple enhancers simultaneously converged on their target gene to synergize in transcriptional initiation, or if such an enhancer network represented the sum of stochastic interactions present in a large cell population. We reasoned that the resolution of this enigma requires the quantitation of chromatin fibre interaction frequencies in small input material. Figure 3A outlines the rationale for this strategy with hypothetical networks being successively reduced in smaller aliquots of input. If such networks represented dynamic and stochastic events the interaction patterns would be expected to display increased variability between the aliquots as the size (n) of a statistical sample affects the standard error for that sample (schematically illustrated in Fig. 3A). Because n is in the denominator of the standard error formula, the standard error decreases as n increases. The smaller the sample, the more variable the responses will be while having more data gives less variation (and more precision) in the results(Kenney 1951). Under this scenario, high biological variability present in very small cell populations would compromise attempts to show reproducibility between such aliquots. To resolve this issue, we prepared three types of samples: One set of ten samples was derived from a large pool of crosslinked and ligated chimeric DNA representing one million cells, each containing 0.88 ng of 3C DNA aliquot corresponding to 176 cells (See methods and Additional file 1: Fig S3A for additional information). The qualities of the libraries are documented in Additional file 1: Figures S3B-E. The other set of nine samples was derived from small cell populations, each corresponding to 177 cells.

**Figure 3:**
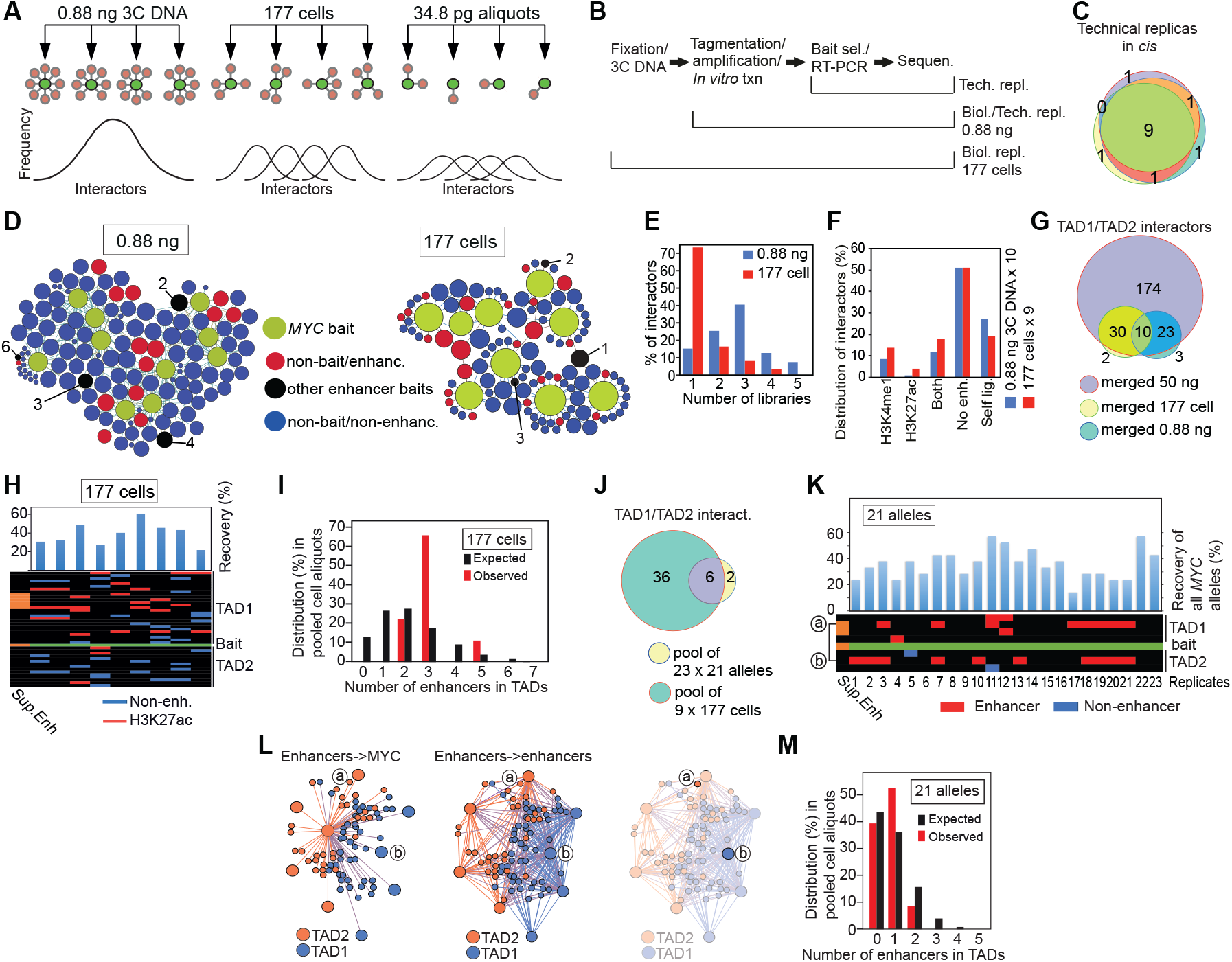
Discrimination between virtual and real enhancer hubs. A) The principle of analysis of stochastic chromatin networks displayed in libraries with lower amount of input. The frequency of the distribution of interactors in the panel showing the 0.88 ng aliquots are predicted to closely follow a normalized distribution profile representative of the initial 3C sample derived from one million cells. Under the assumption that the network represents stochastic interactions, the biological variability is expected to be higher in the 177 cell sample than in the 0.88 ng sample with the highest variability represented by the 34.8 pg/21 alleles. B) Sampling of technical and biological replicas. “Tech.” represents technical whereas “Biol.” represents biological replicas. C) Venn diagram showing the overlap in interactors in *cis* between three different technical replicas. D) Chromatin networks detected by the *MYC* bait on 10 aliquots of 0.88 ng 3C DNA aliquots or 9 different aliquots of 177 cells. The size of each node reflects its connectivity. E) Bar diagram showing the relationship between number of interactors (in *cis*) versus the counts the number of the replicates showing the specific interaction. The bait was omitted from this analysis. F) The stratification of interactors based on the presence or absence of enhancer marks in HCT116 cells. G) Venn diagram demonstrating the overlap between pooled low cell input samples and the 50 ng “ensemble” library. H) Heatmap of enhancer-*MYC* interactions in the flanking TADs within the libraries of the 0.88 ng aliquots. I) The observed frequencies of enhancers impinging on the *MYC* bait compared to the expected frequencies, generated by random resampling of interactors in TADs with enhancer marks impinging on the *MYC* bait scaled from 50 ng 3C DNA input to 177 cells. J) Overlap between pooled libraries from 34.8 pg input and pooled libraries from 177 cells within TADs. K) The overall recovery of *MYC* bait alleles in 23 x 34.8 pg aliquots, each of which corresponds to 21 alleles input material is shown on top of a heatmap of the *MYC* network. The interactors were organized in *cis* keeping their relative position on the physical map. “a” and “b” indicates the location of the interactors represented in L). L) The virtual enhancer hubs observed within the TAD1/2 in the ensemble libraries. The right-most network image identifies lack of physical and direct interaction between nodes “a” and “b”. M) The observed frequencies of enhancers interacting with *MYC* bait compared to the expected frequencies scaled down from 50 ng 3C DNA input to 21 alleles.

Finally, the samples representing the smallest input material consisted of twenty-three aliquots each containing on average 21 alleles (Additional file 1: Fig. S3F), derived from one of the 177-cell samples (Additional file 6: Table S5). To assess the reliability of the Nodewalk technique, we started out by examining the technical variation between RNA libraries generated from 0.88 ng of 3C DNA aliquots, derived from ligated material representing 1 million cells (Fig. 3B). Focusing on interactors in *cis* (Fig. 1G), we observed that the overlap in interactors present in three different aliquots taken from the same initial RNA library, which was prepared from 0.88 ng of 3C DNA (Fig. 3B), generated a technical reproducibility of >90 % (Fig. 3C).

In order to estimate the biological variability among libraries representing independent 177-cell samples (Fig. 3B), we compared the reproducibility of chromatin fibre interactions among the 0.88 ng aliquots to the reproducibility of chromatin fibre interactions among the 177-cell samples, an approach that was simplified by a low variability in the quantity of input and the quality of 3C DNA (Fig. 3D, Additional file 1: Fig. S1A, B and Fig. S3) and a high degree of recovery of the bait alleles (Additional file 1: Fig. S3E). As could be expected (Fig. 3A), >70 % of interactors impinging on the *MYC* bait were detected in only one library among the libraries of the nine 177-cell samples (Fig. 3E,F). In contrast, the libraries of the ten 0.88 ng 3C DNA aliquots showed that >85 % of the interactors were reproduced in two or more libraries. Nonetheless, both types of samples recapitulated the same proportion of interactor categories (Fig. 3F). Importantly, the overlap between the pooled 0.88 ng aliquots or the pooled 177-cell samples and the large ensemble of interactome generated from 3C DNA aliquots corresponding to 10,000 cells exceeded 91%, highlighting that Nodewalk using small input material reliably recapitulated a subset of the interactors already present in the ensemble network (Fig. 3G). As predicted in Figure 3A, the overlap between interactomes present in the libraries that were generated from small input material was only limited (Fig. 3H). We also note that Nodewalk assays of all nine baits from the two TADs in the 0.88 ng 3C DNA aliquots showed little evidence for any enhancer hub interacting with *MYC* (Additional file1: Fig. S3F. See below for more details).

Under the assumption that the enhancer-promoter communications are stochastic, a reduction on cell/allele population sizes should asymptotically reduce the number of enhancers impinging on *MYC* within each aliquot. To assess this issue, we randomly selected interactors from the large ensemble network and scaled it down from 50 ng (corresponding to 10,000 cells) to the level represented by the recovered alleles for the 177 cell aliquots. Figure 3I shows that this approach strongly reduced the expected number of enhancers impinging on *MYC* in such small aliquots. Importantly, there was no significant difference between the expected (generated by 1000 iterations of random sampling from ensemble network) and the actually observed number of enhancers interacting with *MYC* in the libraries derived from the nine 177 cell samples. Although this number is not readily consistent with a static enhancer hub, we further reduced the input sample size to 34.8 pg (Fig. 3G) corresponding to about 21 alleles (Additional file 1: Fig. S3F). For a more robust analysis, only interactors within the *MYC* TADs, which were found to be reproducible across the ensemble network, were retained. This strategy revealed that 6 out of 8 different TAD1/2-specific interactors overlapped entirely with the parental libraries containing a total of 42 TAD1/2-specific interactors, which included non-enhancer as well as enhancer regions in both cases (Fig. 3J). When assessing the distribution of the above-mentioned reproducible interactors among the twenty-three different 34.8 pg aliquots we found that the observed number of different enhancers interacting with *MYC* in each aliquot ranged from 0 to 1, which relatively closely agreed with the permutated number, i.e. between 0 and 3 different enhancers binding *MYC* in each aliquot (Fig. 3K). Note that as there is no interaction between the enhancers labeled as “a” and “b” in the heatmap of Figure 3K within the ensemble network (Fig. 3M), these likely occurred in different cells. Although the number of enhancers impinging on *MYC* increased somewhat when including all *cis* and *trans* enhancers in the ensemble network, Additional file 1: Figure S4 shows that there was still no significant difference between expected and observed number of enhancers impinging on *MYC*. Based on all of these considerations and on the average recovery of the bait being 36.2% (Fig. 3K), we conclude that on average 0.7 enhancer regions per 34.8 ng aliquot interact with on average 7.6 different *MYC* alleles.

Against this conclusion, it could be argued that the Nodewalk technique might have approached a technical limitation for being able to pick up multiple interactions from such small aliquots. However, not only is the number of total alleles recovered in the 23 aliquots exceeding the recovered alleles from each aliquot of the 177 cell samples, but the total number of enhancers within TADs impinging on *MYC* are comparable. Taken together the data strongly implies that the enhancer hubs observed in the ensemble network is only virtual, which is consistent with previous results (Sandhu et al. 2009; Gondor et al. 2010; Ay et al. 2015; Olivares-Chauvet et al. 2016). We conclude, moreover, that although the identity of the involved regions is specific and reproducible in large populations, the dynamics of the interactions represent stochastic events in small populations.

### Potential for interactions versus direct physical interactions and the link to transcriptional activity

To relate interaction frequencies to the frequency of *MYC* transcriptional bursts, we performed RNA FISH analyses using single-stranded probes for introns 1 and 2, followed by DNA FISH analysis (Additional file 1: Fig. S5A,B). Additional file 1: Figure S5C shows that the majority (55.6%) of the *MYC* alleles was transcriptionally active in HCT116 cells, which compares well with the observation above that less than 18% of the *MYC* alleles interact with an enhancer any given time. This result also suggests that the local enhancers do not generally associate with *MYC* once transcription has been initiated. We cannot currently rule out, however, that enhancers might be in physical proximity with transcriptionally active *MYC* alleles without necessarily invoking direct physically contacts. To examine this issue in single cells, we employed the chromatin *in situ* proximity (ChrISP) technique (Chen et al. 2014a; Chen et al. 2014b), which translates the proximity between two different DNA FISH signals into a fluorescent signal only when the epitopes of the differentially labeled DNA FISH probes are <16.2 nm from each other. An 8 kb fragment covering the *MYC* locus and a portion of the super-enhancer (9 kb; Fig. 4A) that showed very frequent interactions with the *MYC* locus (Fig. 4B) and is separated from it by 534 kb (Fig. 4A) were labeled with biotin and digoxygenin, respectively, hybridized to fixed HCT116 cells followed by the visualization of the ChrISP signal (Fig. 4C, Additional file 7: Table S6). Quantitation of the ChrISP signals uncovered that the frequency of super-enhancer-*MYC* proximity is about 50-fold higher than the frequency of direct physical interactions between them as determined by the Nodewalk technique (Fig. 4D). This difference likely reflects that the ChrISP assay measures the ***potential*** for interaction (<16.2 nm), whereas the Nodewalk assay detects only direct physical interactions between interactors, reflecting the length of the formaldehyde monomere (0.59 nm). This result is in agreement with our main conclusion that overt physical interaction between *MYC* and its enhancers is dynamic and stochastic (Fig. 4E) although it also implies that the opportunities for such interactions is manifold higher than observed with the Nodewalk technique.

**Figure 4:**
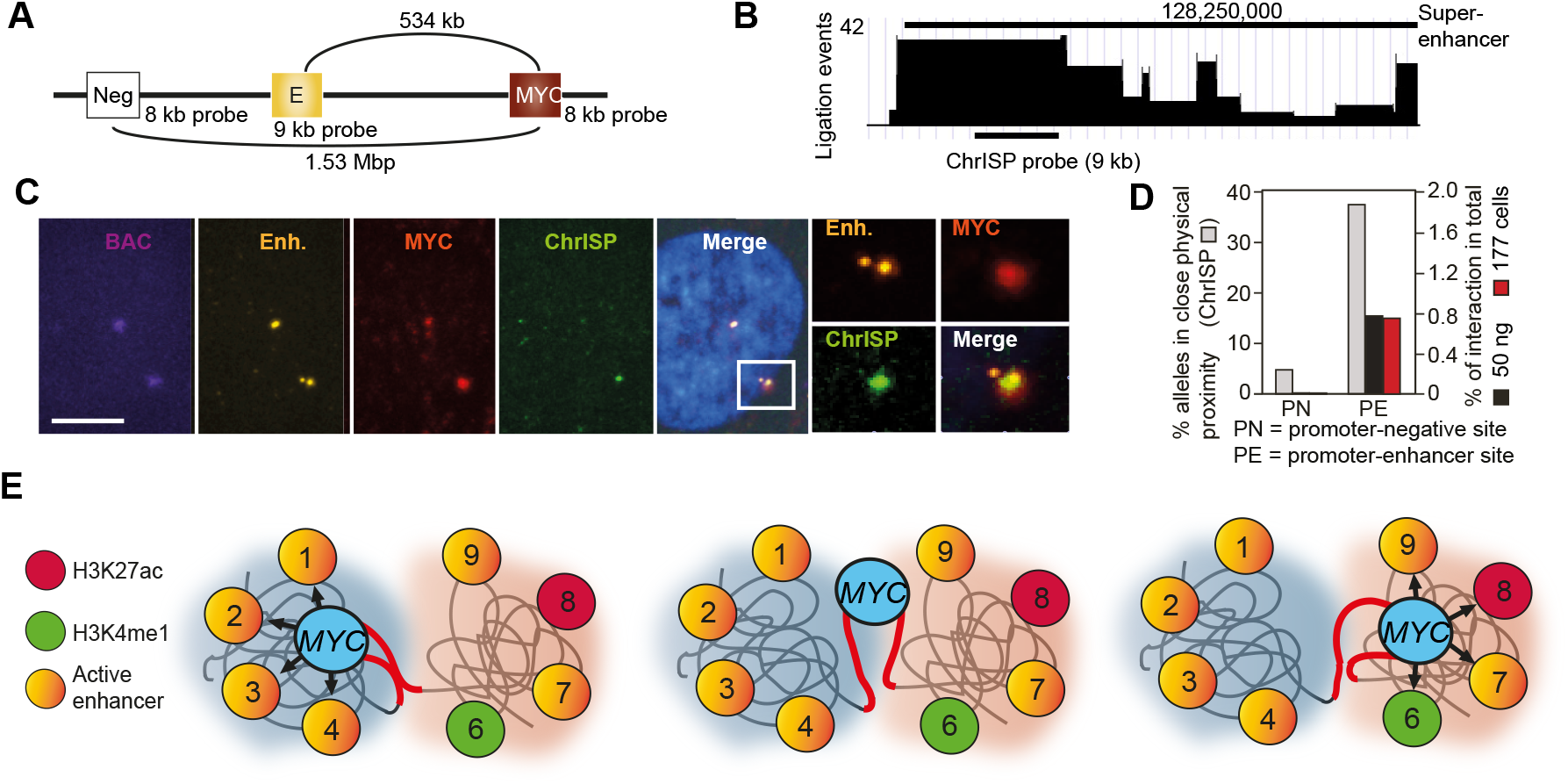
The relationship between direct interactions and potential for interactions. Schematic depiction of the overall (A) and detailed (B) position of the probes used to assess the frequency of proximity between *MYC* and its colorectal super-enhancer in confocal images of single HCT116 cells (C). Bar = 4 μm. Negative controls included omission of secondary antibody and a site not interacting with the *MYC* bait and 1.53 Mbp distal to *MYC* (A). D) The comparison between quantitated frequencies of *MYC*-enhancer proximities with the proportion of the corresponding ratio calculated from the Nodewalk data based on 177 cells or 50 ng. E) Schematic representation of enhancer hubs showing that their likely partner is intra-TAD-specific. The enhancer hubs are postulated to have an increased relative potential for interaction due to compaction of TADs 1 and 2, respectively, in a subset of HCT116 cells with *MYC* hypothesized to be switching between the TADs in mutually exclusive manners.

## Discussion

We have described here the Nodewalk innovation, which uniquely have optimized high resolution with high sensitivity, to assess how variations in 3D chromatin states relate to the transcriptional process in small input materials. In agreement with several other studies (Patrinos et al. 2004; Gavrilov and Razin 2008; Berlivet et al. 2013; Kieffer-Kwon et al. 2013; Dowen et al. 2014; Kim et al. 2014; Liu et al. 2014; Markenscoff-Papadimitriou et al. 2014; Xiang et al. 2014; Ing-Simmons et al. 2015), the chromatin interactome identified from the 50 ng 3C DNA aliquots contains frequent interactions among enhancers that increase the apparent cohesion of the chromatin network. This feature is pronounced especially in cancer cells that contain numerous enhancer regions within the *MYC* TADs. We also show here, however, that this network is only virtual and that the *MYC* gene in reality interacts with an array of distal enhancers distributed in the flanking TADs in a stochastic and hence a largely mutually exclusive manner. This conclusion was possible to make only because of the unique combination of high sensitivity and resolution offered by the Nodewalk technique. By extrapolation, there is thus little evidence for active enhancer hubs simultaneously cooperating to transcriptionally activate *MYC* in single cells. While this conclusion certainly does not rule out the simultaneous presence of multiple enhancer-gene units that aggregate to form transcription factories (Deng et al. 2013), it has the advantage of providing *MYC* with a smorgasbord of distal enhancers. Whether these elements act in synergy or independent from any potential promoter-proximal enhancer function remains to be clarified (Mautner et al. 1995). Irrespective of this consideration, the scenario with numerous distal enhancers conceivably serves to render *MYC* expression and hence cancer cells more adaptive to fluctuating environments experienced during the neoplastic process (Mautner et al. 1995). The opposite interpretation, invoking the possibility that the enhancers continuously interact to boost *MYC* transcription would require static chromatin fibre interactions, which is not consistent with the dynamic juxtapositions between the CRC enhancer and *MYC*.

To reach these conclusions, we optimized the Nodewalk technique to yield a sensitivity of at least 21 alleles, corresponding to 7 cells, and yet keeping a high resolution of interacting chromatin fragments. However, by pushing the boundaries for such a sensitivity, we faced a conundrum: How to discriminate biological variability in chromatin fibre interactomes from technical variation among chromatin networks generated from unique, small cell populations? We used three strategies to deal with this issue. First, we have shown that the technical variation of the Nodewalk analysis of small aliquots from the same linearly amplified RNA library (i.e. representing the same initial chimeric DNA library) is very low (<12%). Second, we observed extensive overlap in the identity of the interactors between the pool of twenty-three 34.8pg 3C DNA aliquots (corresponding to 21 alleles) or the nine 177-cell samples and the ensemble library generated from two 50 ng 3C DNA aliquots (derived from one million cells), further demonstrating the reproducibility of the assay in small input material. Third, by binning the interactome data generated from small samples in successively larger windows, the overall patterns of the ensemble network could be reproduced to demonstrate the stochastic but preferential patterns of interactions between *MYC* and its enhancers (Additional file 1: Fig. S6). Moreover, a comparison between observed and expected data generated by permutation analyses uncovered that interactions between *MYC* and its enhancers display stochastic features in small cell populations. While it could be argued that limitations in the ligation step would preclude identification of multiple enhancers interacting with *MYC* in small samples, such as the 34.8 pg aliquots, we note that the number of enhancers impinging on *MYC* does not increase even when pooling the recovery of all 23 of the 34.8 pg aliquots to yield a total of 174 alleles. Taken together, we argue that the Nodewalk technique represent a significant improvement over competing techniques to reliably discriminate between real and virtual chromatin networks.

Our data also addresses whether it is the enhancer or the gene driving the interaction patterns. Given that the interactions between the TADs are generally infrequent and that *MYC* taps equally well into the two neighboring TADs both in HCT116 cells and HCECs, we submit that the mobility of the inter-TAD boundary containing *MYC* drives *MYC*-enhancer communications. This interpretation is underscored by the ChrISP analysis showing that a significant proportion of the *MYC* alleles are in relatively close physical proximity to the colorectal super-enhancer within TAD1, suggesting the existence of a more compacted TAD conformation in a subset of the cells to increase the potential for interactions. Thus, 43% of all TAD1-specific interactors distribute over a large 1 Mbp region represented by TAD1 (Fig. 2F), despite that 37% of all alleles at the same time display close physical proximity between a small portion of the super-enhancer and *MYC* (Fig. 4D) From this it follows that a large fraction of the *MYC* alleles is proximal to the entire TAD while being physically juxtaposed to the colorectal super-enhancer. Such a process might place enhancer hubs in relative physical proximity to each other although the frequency of direct physical interactions between them is likely dynamic and hence not detectable by Nodewalk in small cell populations. With *MYC* strategically placed between the two flanking TADs, we further propose that such a position enhances the ability of *MYC* to stochastically engage enhancers from either TAD (Fig. 4E). This reasoning is in keeping with the observation that *MYC* has the highest dynamic importance index in comparison with enhancers positioned within either of the TADs in the virtual network. We conclude that while *MYC* generally interacts with only one enhancer at a time in very small sample inputs, colon cancer cells have organized a chromatin environment in which enhancers might be in relative proximity to each other in a TAD-specific manner to facilitate *MYC*-enhancer communications. The plasticity underlying this process provides the cancer cell with a selective advantage as multiple signaling pathways converge on different sets of *MYC* enhancers in cancer cells (Hnisz et al. 2015). Combined with the flexibility incurred by being positioned at an inter-TAD boundary, such an accessibility of enhancers might ensure *MYC* activation to drive excessive cell proliferation irrespective of a changing microenvironment that the cancer cell might experience during the neoplastic process.

To sum up, by applying the Nodewalk technique to very small cell populations, we have been able to document that the chromatin networks within the two TADs impinging on *MYC* have evolved to facilitate redundant mechanisms of *MYC* activation in cancer cells. The major advantage of this decentralized network topology is that there might be no “single point of failure” within the network to ensure the potential for continuous activation of *MYC* to increase the fitness of cancer cells by promoting their adaptability to changing microenvironments. Such a network structure might necessitate the identification of therapeutic strategies that focus on the inter-TAD boundary rather than the super-enhancers to attenuate *MYC* expression in cancer patients. On a more general note, the extreme sensitivity and versatility of the Nodewalk technique has opened up numerous, previously inaccessible applications, such as the deciphering of cellular heterogeneity in chromatin structures within solid and liquid tumor biopsies and circulating tumor cells, where subpopulations of cells are identified by marker antibodies. Such an approach might also be essential to determine the precise order of events underlying stochastic transcriptional activation driving cancer evolution.

## Acknowledgements

The authors would like to acknowledge support from Science for Life Laboratory, the National Genomics Infrastructure, NGI, and Uppmax for providing assistance in massive parallel sequencing and computational infrastructure as well as the extensive data sets from ENCODE. This work was supported by the Swedish Cancer Foundation (CAN2016/576 (AG); CAN2016/616 (JPS), VR-NT (2013-4511 (RO)), VR-M (2015-02312 (PS); 2014-3683 (RO); 2016-03108 (AG)), Karolinska Institutet (AG, RO), Åke Wiberg Stiftelse (M16-0090; AG)), Swedish Pediatric Cancer Foundation (2015-0129; AG)), and the KA Wallenberg Foundation (Clinical Epigenetics; RO).

## Availability of data and materials

All the sequence data have been deposited to the NCBI Gene Expression Omnibus (https://www.ncbi.nlm.nih.gov/geo/) under the accession number of GSE76049.

## Legends to the supplemental figures

**Figure S1:**
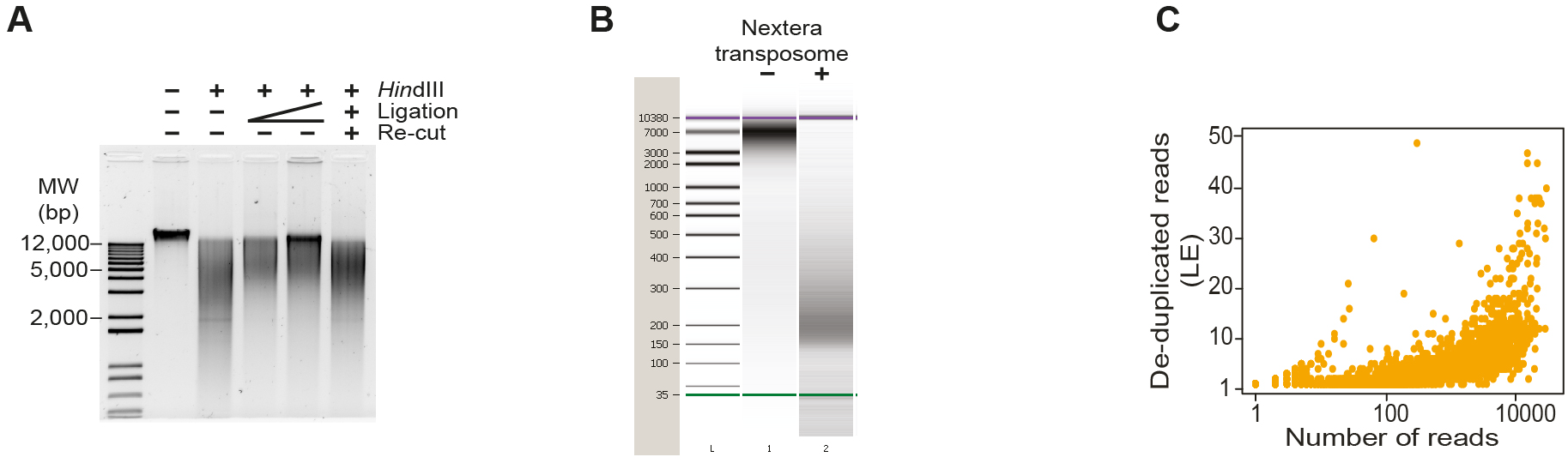
A) Digestion and ligation efficiencies analyzed by agarose gel electrophoresis. B) Nextera tagmentation fragments 3C DNA to uniform sizes. C) Comparison between number of raw reads and the de-duplicated reads (LE).

**Figure S2:**
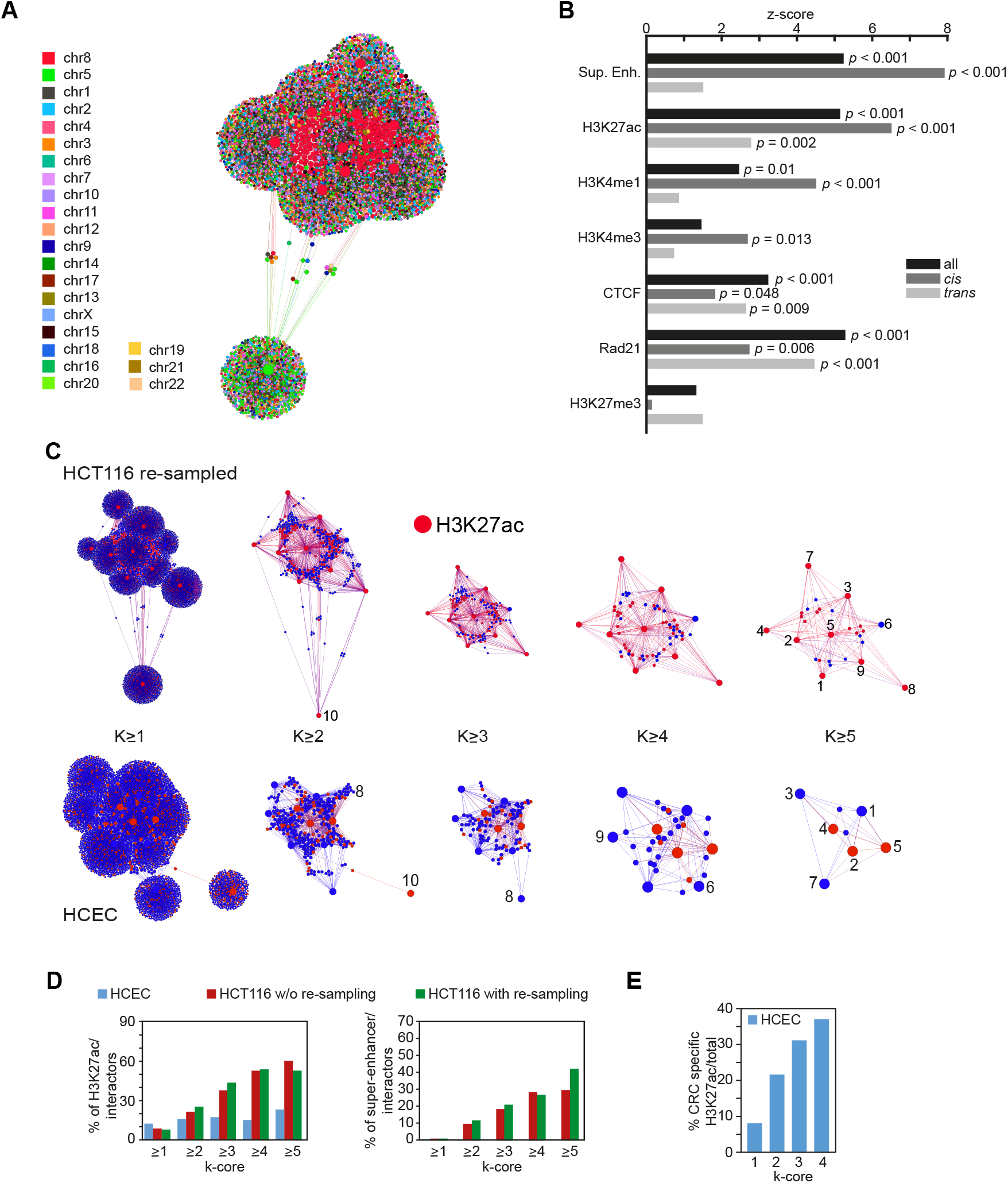
Enrichment of enhancer chromatin hubs identified by Nodewalk. A) Map of inter-chromosomal interactome generated from 10 different baits with each chromosome color-coded as indicated in the image. B) Enrichment analysis of chromatin marks at the network nodes. P-values were estimated from 1,000 permutations (see the methods). C) The re-sampled network structure from HCT116 and HCEC stratified by their k-core values. The nodes of the red identify regions overlapping H3K27ac peaks. The size of each node reflects the number of the interactors. D) The comparison of the proportion of either H3k27ac peaks (left) or super-enhancer (right) between HCEC, HCT116 and re-sampled HCT116 stratified with k-core values. E) The enrichment of cancer-specific H3K27ac mark on HCEC interactors.

**Figure S3:**
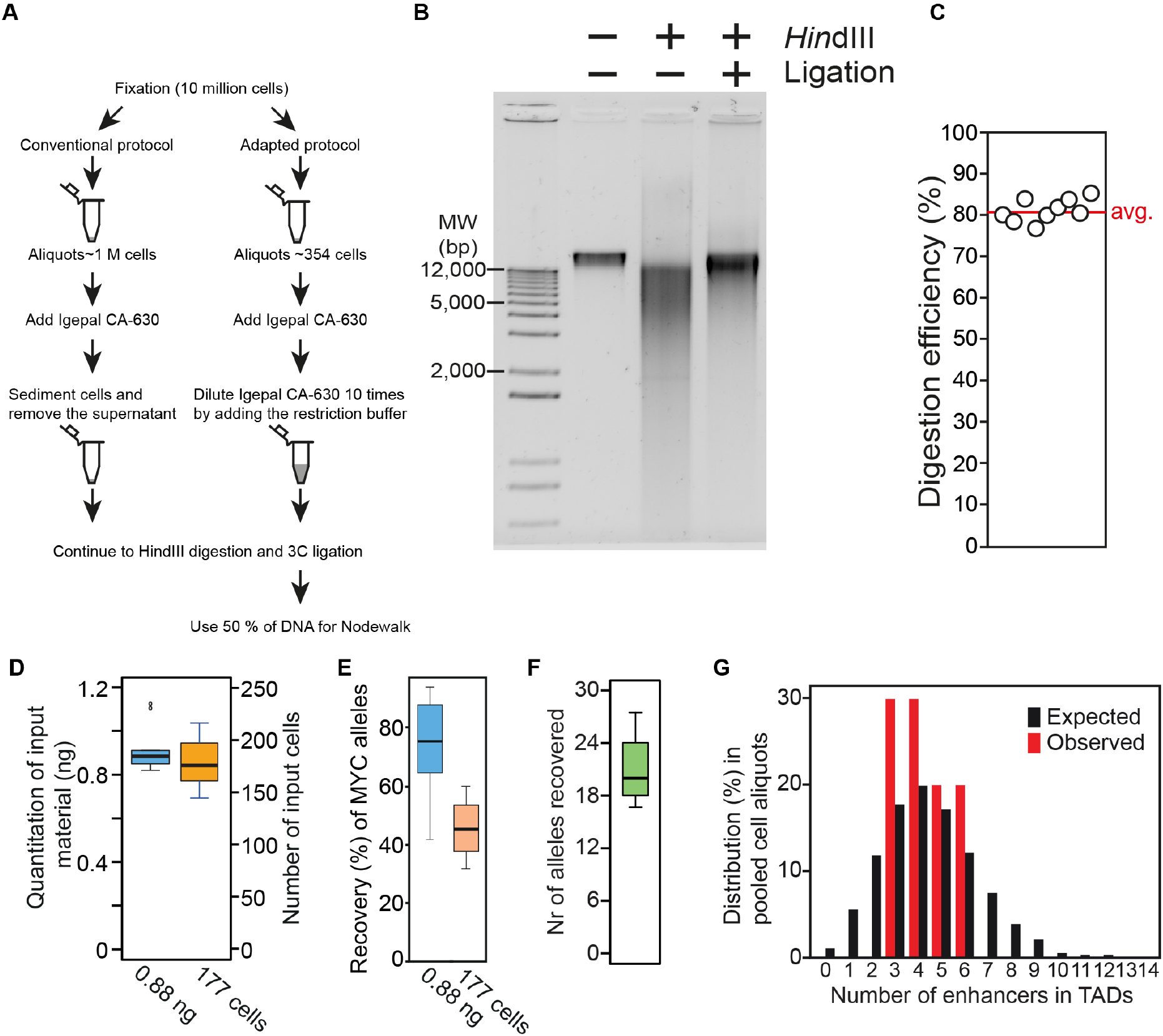
The modified Nodewalk protocol adapted for lower amount of the cells. A) Flow scheme. B) Visualization of Hind III digestion and 3C ligation using adapted protocol with 10,000 cells. C) Digestion efficiency of chromatin DNA (at the *MYC* bait) of 9 different aliquots of input material corresponding to 177 cells. D) The quantification of input DNA from 0.88 ng aliquots of 3C DNA or 177 cells. E) Recovery of *MYC* alleles in chromatin network generated by Nodewalk analyses of individual aliquots as indicated. F) The quantification of input from 34.8 pg of 3C DNA. G) The observed frequencies of enhancers impinging on the MYC bait compared to the expected frequencies, generated by random resampling of interactors in TADs scaled from 50 ng 3C DNA input to 0.88 ng.

**Figure S4:**
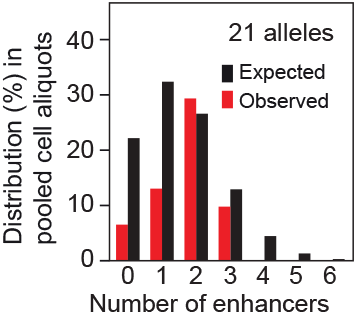
The observed frequencies of enhancers interacting with MYC bait compared to the expected frequencies scaled down from 50 ng 3C DNA input to 21 alleles. Values were calculated from the interactors in whole genome

**Figure S5:**
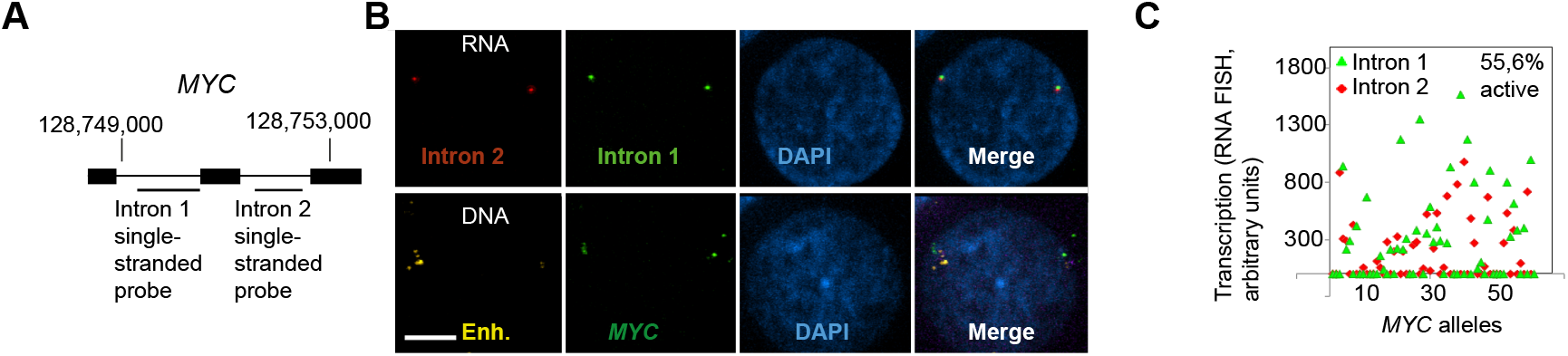
RNA FISH analysis of *MYC* transcription. A) The position of the single-stranded intron probes used for RNA FISH analysis. B) Sequential confocal RNA/DNA FISH images exemplifying the identification of active and inactive MYC alleles, respectively. Bar= 4 μm. C) Summary of the RNA FISH signals (green=intron 1; red=intron 2) to depict ongoing MYC transcription in HCT116 cells. The various proportions of the red and green signals in the chart likely reflect partial transcripts and/or partial RNA processing.

**Figure S6:**
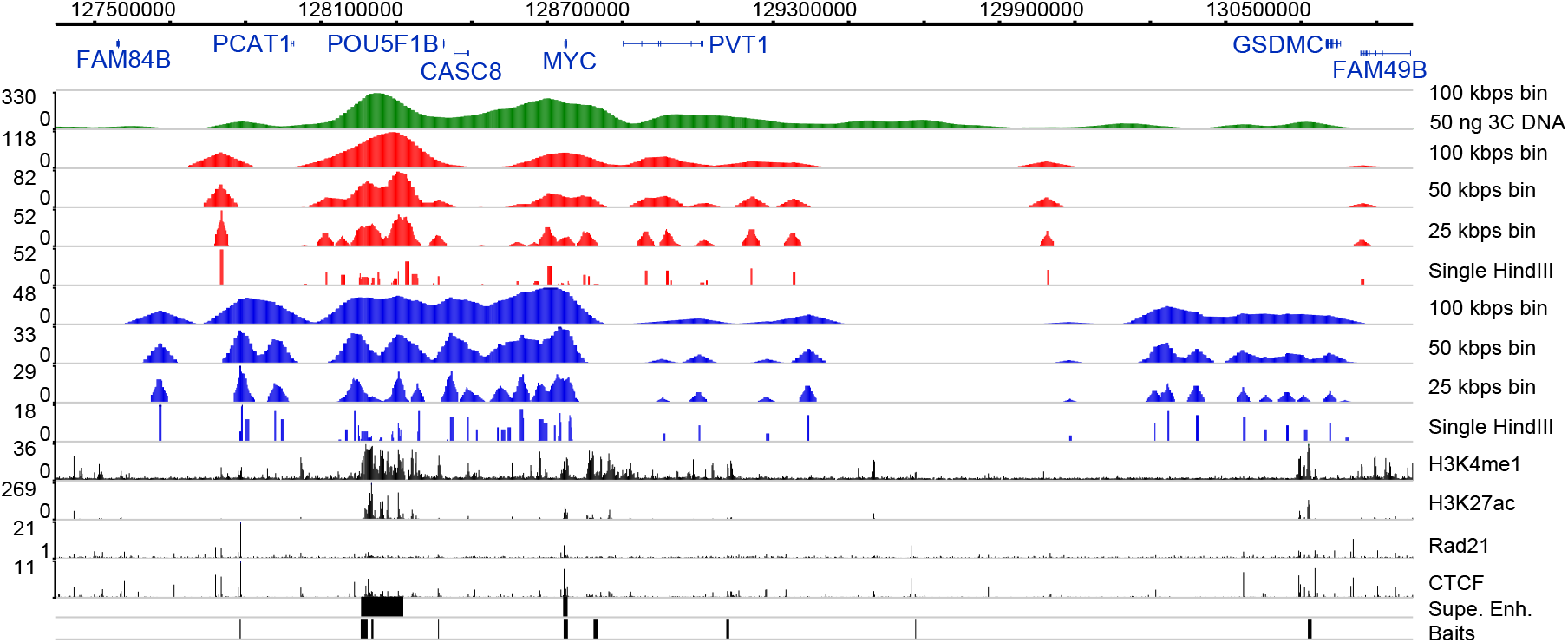
The stochastic character of the *MYC* network. Comparison of interaction profiles in the TADs flanking MYC with interaction profiles binned into larger windows resulting from Nodewalk analyses using 50 ng 3C DNA/10,000 cells input (green), 0.88 ng 3C DNA aliquots (red) and 177 cell aliquots (blue).

## References

Ay F, Vu TH, Zeitz MJ, Varoquaux N, Carette JE, Vert JP, Hoffman AR, Noble WS. 2015. Identifying multi-locus chromatin contacts in human cells using tethered multiple 3C. BMC genomics 16: 121.

Bastian M. HS, Jacomy M. 2009. Gephi: an open source software for exploring and manipulating networks. International AAAI Conference on Weblogs and Social Media.

Berlivet S, Paquette D, Dumouchel A, Langlais D, Dostie J, Kmita M. 2013. Clustering of tissue-specific sub-TADs accompanies the regulation of HoxA genes in developing limbs. PLoS Genet 9: e1004018.

Chen X, Shi C, Yammine S, Gondor A, Ronnlund D, Fernandez-Woodbridge A, Sumida N, Widengren J, Ohlsson R. 2014a. Chromatin in situ proximity (ChrISP): Single-cell analysis of chromatin proximities at a high resolution. Biotechniques 56: 117–124.

Chen X, Yammine S, Shi C, Tark-Dame M, Gondor A, Ohlsson R. 2014b. The visualization of large organized chromatin domains enriched in the H3K9me2 mark within a single chromosome in a single cell. Epigenetics 9: 1439–1445.

Csardi G, Nepusz, T.,. 2006. The igraph software package for complex network research. InterJournal, Complex Systems: 1695.

Dekker J, Rippe K, Dekker M, Kleckner N. 2002. Capturing chromosome conformation. Science 295: 1306–1311.

Deng B, Melnik S, Cook PR. 2013. Transcription factories, chromatin loops, and the dysregulation of gene expression in malignancy. Seminars in cancer biology 23: 65–71.

Dixon JR, Selvaraj S, Yue F, Kim A, Li Y, Shen Y, Hu M, Liu JS, Ren B. 2012. Topological domains in mammalian genomes identified by analysis of chromatin interactions. Nature 485: 376–380.

Dowen JM, Fan ZP, Hnisz D, Ren G, Abraham BJ, Zhang LN, Weintraub AS, Schuijers J, Lee TI, Zhao K et al. 2014. Control of cell identity genes occurs in insulated neighborhoods in mammalian chromosomes. Cell 159: 374–387.

Dryden NH, Broome LR, Dudbridge F, Johnson N, Orr N, Schoenfelder S, Nagano T, Andrews S, Wingett S, Kozarewa I et al. 2014. Unbiased analysis of potential targets of breast cancer susceptibility loci by Capture Hi-C. Genome Res 24: 1854–1868.

Du Z, Zheng H, Huang B, Ma R, Wu J, Zhang X, He J, Xiang Y, Wang Q, Li Y et al. 2017. Allelic reprogramming of 3D chromatin architecture during early mammalian development. Nature 547: 232–235.

Fullwood MJ, Liu MH, Pan YF, Liu J, Xu H, Mohamed YB, Orlov YL, Velkov S, Ho A, Mei PH et al. 2009. An oestrogen-receptor-alpha-bound human chromatin interactome. Nature 462: 58–64.

Gavrilov AA, Razin SV. 2008. Spatial configuration of the chicken alpha-globin gene domain: immature and active chromatin hubs. Nucleic Acids Res 36: 4629–4640.

Gondor A, Rougier C, Ohlsson R. 2008. High-resolution circular chromosome conformation capture assay. Nat Protoc 3: 303–313.

Gondor A, Woodbridge AF, Shi C, Aurell E, Imreh M, Ohlsson R. 2010. Window into the complexities of chromosome interactomes. Cold Spring Harb Symp Quant Biol 75: 493–500.

Hager GL, McNally JG, Misteli T. 2009. Transcription dynamics. Molecular cell 35: 741–753.

Hay D, Hughes JR, Babbs C, Davies JO, Graham BJ, Hanssen LL, Kassouf MT, Oudelaar AM, Sharpe JA, Suciu MC et al. 2016. Genetic dissection of the alpha-globin super-enhancer in vivo. Nature genetics 48: 895–903.

Hnisz D, Abraham BJ, Lee TI, Lau A, Saint-Andre V, Sigova AA, Hoke HA, Young RA. 2013. Super-enhancers in the control of cell identity and disease. Cell 155: 934–947.

Hnisz D, Schuijers J, Lin CY, Weintraub AS, Abraham BJ, Lee TI, Bradner JE, Young RA. 2015. Convergence of developmental and oncogenic signaling pathways at transcriptional super-enhancers. Mol Cell 58: 362–370.

Hughes JR, Roberts N, McGowan S, Hay D, Giannoulatou E, Lynch M, De Gobbi M, Taylor S, Gibbons R, Higgs DR. 2014. Analysis of hundreds of cis-regulatory landscapes at high resolution in a single, high-throughput experiment. Nature genetics 46: 205–212.

Ing-Simmons E, Seitan VC, Faure AJ, Flicek P, Carroll T, Dekker J, Fisher AG, Lenhard B, Merkenschlager M. 2015. Spatial enhancer clustering and regulation of enhancer-proximal genes by cohesin. Genome Res 25: 504–513.

Ji X, Dadon DB, Powell BE, Fan ZP, Borges-Rivera D, Shachar S, Weintraub AS, Hnisz D, Pegoraro G, Lee TI et al. 2015. 3D Chromosome Regulatory Landscape of Human Pluripotent Cells. Cell stem cell doi:10.1016/j.stem.2015.11.007.

Kenney JF. 1951. Mathematics of Statistics,. Chapman & Hall LTD.

Kieffer-Kwon KR, Tang Z, Mathe E, Qian J, Sung MH, Li G, Resch W, Baek S, Pruett N, Grontved L et al. 2013. Interactome maps of mouse gene regulatory domains reveal basic principles of transcriptional regulation. Cell 155: 1507–1520.

Kim T, Cui R, Jeon YJ, Lee JH, Lee JH, Sim H, Park JK, Fadda P, Tili E, Nakanishi H et al. 2014. Long-range interaction and correlation between MYC enhancer and oncogenic long noncoding RNA CARLo-5. Proc Natl Acad Sci U S A 111: 4173–4178.

Langer S, Geigl JB, Ehnle S, Gangnus R, Speicher MR. 2005. Live cell catapulting and recultivation does not change the karyotype of HCT116 tumor cells. Cancer Genet Cytogenet 161: 174–177.

Li G, Ruan X, Auerbach RK, Sandhu KS, Zheng M, Wang P, Poh HM, Goh Y, Lim J, Zhang J et al. 2012. Extensive promoter-centered chromatin interactions provide a topological basis for transcription regulation. Cell 148: 84–98.

Lieberman-Aiden E, van Berkum NL, Williams L, Imakaev M, Ragoczy T, Telling A, Amit I, Lajoie BR, Sabo PJ, Dorschner MO et al. 2009. Comprehensive mapping of long-range interactions reveals folding principles of the human genome. Science 326: 289–293.

Liu Z, Legant WR, Chen BC, Li L, Grimm JB, Lavis LD, Betzig E, Tjian R. 2014. 3D imaging of Sox2 enhancer clusters in embryonic stem cells. Elife 3: e04236.

Lohmann G, Margulies DS, Horstmann A, Pleger B, Lepsien J, Goldhahn D, Schloegl H, Stumvoll M, Villringer A, Turner R. 2010. Eigenvector centrality mapping for analyzing connectivity patterns in fMRI data of the human brain. PloS one 5: e10232.

Loven J, Hoke HA, Lin CY, Lau A, Orlando DA, Vakoc CR, Bradner JE, Lee TI, Young RA. 2013. Selective inhibition of tumor oncogenes by disruption of super-enhancers. Cell 153: 320–334.

Markenscoff-Papadimitriou E, Allen WE, Colquitt BM, Goh T, Murphy KK, Monahan K, Mosley CP, Ahituv N, Lomvardas S. 2014. Enhancer interaction networks as a means for singular olfactory receptor expression. Cell 159: 543–557.

Mautner J, Joos S, Werner T, Eick D, Bornkamm GW, Polack A. 1995. Identification of two enhancer elements downstream of the human c-myc gene. Nucleic Acids Res 23: 72–80.

Nagano T, Lubling Y, Stevens TJ, Schoenfelder S, Yaffe E, Dean W, Laue ED, Tanay A, Fraser P. 2013. Single-cell Hi-C reveals cell-to-cell variability in chromosome structure. Nature 502: 59–64.

Naumova N, Smith EM, Zhan Y, Dekker J. 2012. Analysis of long-range chromatin interactions using Chromosome Conformation Capture. Methods 58: 192–203.

Olivares-Chauvet P, Mukamel Z, Lifshitz A, Schwartzman O, Elkayam NO, Lubling Y, Deikus G, Sebra RP, Tanay A. 2016. Capturing pairwise and multi-way chromosomal conformations using chromosomal walks. Nature 540: 296–300.

Patrinos GP, de Krom M, de Boer E, Langeveld A, Imam AM, Strouboulis J, de Laat W, Grosveld FG. 2004. Multiple interactions between regulatory regions are required to stabilize an active chromatin hub. Genes Dev 18: 1495–1509.

Picelli S, Bjorklund AK, Reinius B, Sagasser S, Winberg G, Sandberg R. 2014. Tn5 transposase and tagmentation procedures for massively scaled sequencing projects. Genome research 24: 2033–2040.

Rao SS, Huntley MH, Durand NC, Stamenova EK, Bochkov ID, Robinson JT, Sanborn AL, Machol I, Omer AD, Lander ES et al. 2014. A 3D map of the human genome at kilobase resolution reveals principles of chromatin looping. Cell 159: 1665–1680.

Sanchez A, Golding I. 2013. Genetic determinants and cellular constraints in noisy gene expression. Science 342: 1188–1193.

Sandhu KS, Shi C, Sjolinder M, Zhao Z, Gondor A, Liu L, Tiwari VK, Guibert S, Emilsson L, Imreh MP et al. 2009. Nonallelic transvection of multiple imprinted loci is organized by the H19 imprinting control region during germline development. Genes Dev 23: 2598–2603.

Schwartzman O, Mukamel Z, Oded-Elkayam N, Olivares-Chauvet P, Lubling Y, Landan G, Izraeli S, Tanay A. 2016. UMI-4C for quantitative and targeted chromosomal contact profiling. Nature methods 13: 685–691.

Seidman SB. 1983. Network structure and minimum degree. Social Networks 5: 269–287.

Shi J, Whyte WA, Zepeda-Mendoza CJ, Milazzo JP, Shen C, Roe JS, Minder JL, Mercan F, Wang E, Eckersley-Maslin MA et al. 2013. Role of SWI/SNF in acute leukemia maintenance and enhancer-mediated Myc regulation. Genes & development 27: 2648–2662.

Stevens TJ, Lando D, Basu S, Atkinson LP, Cao Y, Lee SF, Leeb M, Wohlfahrt KJ, Boucher W, O’Shaughnessy-Kirwan A et al. 2017. 3D structures of individual mammalian genomes studied by single-cell Hi-C. Nature 544: 59–64.

Williams RL, Jr., Starmer J, Mugford JW, Calabrese JM, Mieczkowski P, Yee D, Magnuson T. 2014. fourSig: a method for determining chromosomal interactions in 4C-Seq data. Nucleic Acids Res 42: e68.

Xiang JF, Yin QF, Chen T, Zhang Y, Zhang XO, Wu Z, Zhang S, Wang HB, Ge J, Lu X et al. 2014. Human colorectal cancer-specific CCAT1-L lncRNA regulates long-range chromatin interactions at the MYC locus. Cell Res 24: 513–531.

Zhao H, Sifakis EG, Sumida N, Millan-Arino L, Scholz BA, Svensson JP, Chen X, Ronnegren AL, Mallet de Lima CD, Varnoosfaderani FS et al. 2015. PARP1- and CTCF-Mediated Interactions between Active and Repressed Chromatin at the Lamina Promote Oscillating Transcription. Mol Cell 59: 984–997.

Zhao Z, Tavoosidana G, Sjolinder M, Gondor A, Mariano P, Wang S, Kanduri C, Lezcano M, Sandhu KS, Singh U et al. 2006. Circular chromosome conformation capture (4C) uncovers extensive networks of epigenetically regulated intra- and interchromosomal interactions. Nat Genet 38: 1341–1347.

Zhou X, Maricque B, Xie M, Li D, Sundaram V, Martin EA, Koebbe BC, Nielsen C, Hirst M, Farnham P et al. 2011. The Human Epigenome Browser at Washington University. Nature methods 8: 989–990.

